# Measles skin rash: infection of lymphoid and myeloid cells in the dermis precedes viral dissemination to keratinocytes in the epidermis

**DOI:** 10.1101/867531

**Authors:** Brigitta M. Laksono, Paola Fortugno, Bernadien M. Nijmeijer, Rory D. de Vries, Sonia Cordisco, Stefan van Nieuwkoop, Thijs Kuiken, Teunis B. H. Geijtenbeek, W. Paul Duprex, Francesco Brancati, Rik L. de Swart

**Author notes:** Department of Life, Health and Environmental Sciences, University of L’Aquila, L’Aquila, Italy (Sonia Cordisco, Francesco Brancati). Correspondence to: Dr. Rik L. de Swart, Department of Viroscience, Erasmus MC. Wytemaweg 80, 3015 CN Rotterdam, The Netherlands.; E.

## Abstract

Measles is characterised by fever and a maculopapular skin rash, which is associated with immune clearance of measles virus (MV)-infected cells. Histopathological analyses of skin biopsies from humans and non-human primates (NHPs) with measles rash have identified MV-infected keratinocytes and mononuclear cells in the epidermis, around hair follicles and near sebaceous glands. Here, we address the pathogenesis of measles skin rash by combining data from experimentally infected NHPs, *ex vivo* infection of human skin sheets and *in vitro* infection of primary human keratinocytes. Longitudinal analysis of the skin of experimentally MV-infected NHPs demonstrated that infection in the skin precedes onset of rash by several days. MV infection was initiated in lymphoid and myeloid cells in the dermis before dissemination to the epidermal keratinocytes. These data were in good concordance with *ex vivo* MV infections of human skin sheets, in which dermal cells were more targeted than the epidermal ones. To address viral dissemination to the epidermis and to determine whether the dissemination is receptor-dependent, we performed experimental infections of primary keratinocytes collected from healthy or nectin-4-deficient donors. These experiments demonstrated that MV infection of keratinocytes is nectin-4-dependent, and nectin-4 expression was higher in differentiated than in proliferating keratinocytes. Based on these data, we hypothesise that measles skin rash is initiated by migrating MV-infected lymphocytes that infect dermal skin-resident CD150^+^ immune cells. The infection is subsequently disseminated from the dermal papillae to nectin-4^+^ keratinocytes in the basal epidermis. Lateral spread of MV infection is observed in the superficial epidermis, most likely due to the higher level of nectin-4 expression on differentiated keratinocytes. Finally, MV-infected cells are cleared by infiltrating immune cells, causing hyperaemia and oedema, which give the appearance of morbilliform skin rash.

**Author Summary:** Several viral infections are associated with skin rash, including parvovirus B19, human herpesvirus type 6, dengue virus and rubella virus. However, the archetype virus infection that leads to skin rash is measles. Although all of these viral exanthemata often appear similar, their pathogenesis is different. In the case of measles, the appearance of skin rash is a sign that the immune system is clearing MV-infected cells from the skin. How the virus reaches the skin and is locally disseminated remains unknown. Here we combine observations and expertise from pathologists, dermatologists, virologists and immunologists to delineate the pathogenesis of measles skin rash. We show that MV infection of dermal myeloid and lymphoid cells precedes viral dissemination to the epidermal keratinocytes. We speculate that immune-mediated clearance of these infected cells results in hyperaemia and oedema, explaining the redness of the skin and the slightly elevated spots of the morbilliform rash.

## Introduction

Measles virus (MV) is a highly contagious enveloped virus with a negative single-stranded RNA genome, that belongs to the family *Paramyxoviridae*, genus *Morbillivirus* (PMID: 31609197). Measles is associated with fever, cough and a characteristic maculopapular skin rash [1]. MV utilises two cellular receptors to infect its target cells: CD150 and nectin-4 [2–4]. CD150 plays a crucial role during viral entry and systemic dissemination. It is expressed on subsets of immune cells, including macrophages, dendritic cells (DCs) and lymphocytes. Nectin-4 is crucial for viral transmission to the next host. It is an adherens junction protein expressed at the basolateral surface of differentiated respiratory epithelial cells and is involved in the maintenance of epithelium integrity [5, 6].

Following entry of MV into the respiratory tract, the primary infection of myeloid cells leads to a cell-associated viremia mediated by CD150^+^ lymphocytes, resulting in systemic disease [7–9]. During a clinically silent incubation phase of 7 to 10 days, circulating MV-infected lymphocytes migrate into various tissues and transmit the virus to susceptible tissue-resident CD150^+^ immune cells and nectin-4^+^ epithelial cells. Basolateral infection of respiratory epithelial cells leads to the apical release of nascent virions into the lumen of the respiratory tract [10–12]. Shedding is associated with the onset of prodromal clinical signs such as fever and cough [9, 13]. Maculopapular skin rash and conjunctivitis follow a few days later [9] and are associated with onset of MV-specific cellular immune responses [13]. Patients with a compromised cellular immune system do not develop rash or conjunctivitis, but are at high risk of developing severe disease [14].

In histopathological studies of human skin biopsies, measles skin rash is mostly characterised by infection and necrosis of keratinocytes and mononuclear cells in the epidermis, and multinucleated giant cells located in proximity to hair follicles and sebaceous glands [15, 16]. It has been postulated that measles rash starts by infection of dermal endothelial cells [17]. However, these cells neither express CD150 nor nectin-4 [18, 19]. Moreover, we have previously identified MV-infected lymphocytes and DCs in the skin of experimentally infected non-human primates preceding onset of skin rash [8]. Besides CD150^+^ and nectin-4^+^ cells, other cells that express DC-SIGN or Langerin could play a role in the pathogenesis of measles skin rash, since DC-SIGN and Langerin facilitate attachment, but not entry, of MV and thus potentially help in spreading the infection in the skin.

In order to understand the pathogenesis of measles skin rash, it is important to understand both the architecture of the skin and the spatial organisation of cell subsets that express either CD150 or nectin-4. The dermis is vascularised and contains several subsets of immune cells that express CD150. These include a network of myeloid DCs and clusters of tissue-resident CD4^+^ and CD8^+^ T cells [20–22]. In contrast to the dermis, the epidermis is not vascularised. It mainly consists of keratinocytes, with an interdigitating network of Langerhans cells (LCs) and melanocytes [23]. The epidermis comprises of proliferating keratinocytes at the basal lamina that differentiate towards the skin surface. Keratinocytes express nectin-4 and expression levels increase during differentiation. It is known that keratinocytes are susceptible to MV infection [24]. The top layer of the epidermis, the stratum corneum, consists of a layer of dead keratinocytes called corneocytes. Interestingly, immune cells and nutrients can only reach the epidermis by migration and diffusion, respectively, from the superficial dermis through the basal lamina. The pilosebaceous unit begins at the epidermis and extends into the dermis, where the surrounding tissue is usually better vascularised. Therefore, tissue-resident lymphocytes are often seen in close association with these structures [21]. Hair follicles are mainly constituted of keratinocytes that express high levels of nectin-4 explaining their propensity for MV infection [25].

During viraemia, systemic dissemination of MV is mediated by circulating MV-infected CD150^+^ lymphocytes. However, how these cells infiltrate the skin, ultimately resulting in skin rash, remains largely unknown. In this study we aimed to identify the cells involved in MV infection of the skin and to understand the pathogenesis of skin rash. We demonstrate that MV infection of lymphoid and myeloid cells in the superficial dermis precedes dissemination to epidermal keratinocytes, which is followed by onset of the typical skin rash.

## Results

### MV skin infection precedes onset of rash in experimentally-infected non-human primates (NHPs)

We retrospectively analysed data from cynomolgus macaques (*Macaca fascicularis*) inoculated with recombinant MV (rMV) strains expressing enhanced green fluorescent protein (EGFP) [26]. Fluorescent spots, indicating the presence of MV-infected cells, became detectable in the skin around 8 days post-inoculation (dpi), although skin rash only became prominent between 11 and 13 dpi (Fig 1) [27]. We previously reported that by 9 dpi, *i.e*. before the onset of rash, infected cells in the skin mainly consisted of EGFP^+^ lymphocytes and DCs [8].

**Fig 1.**
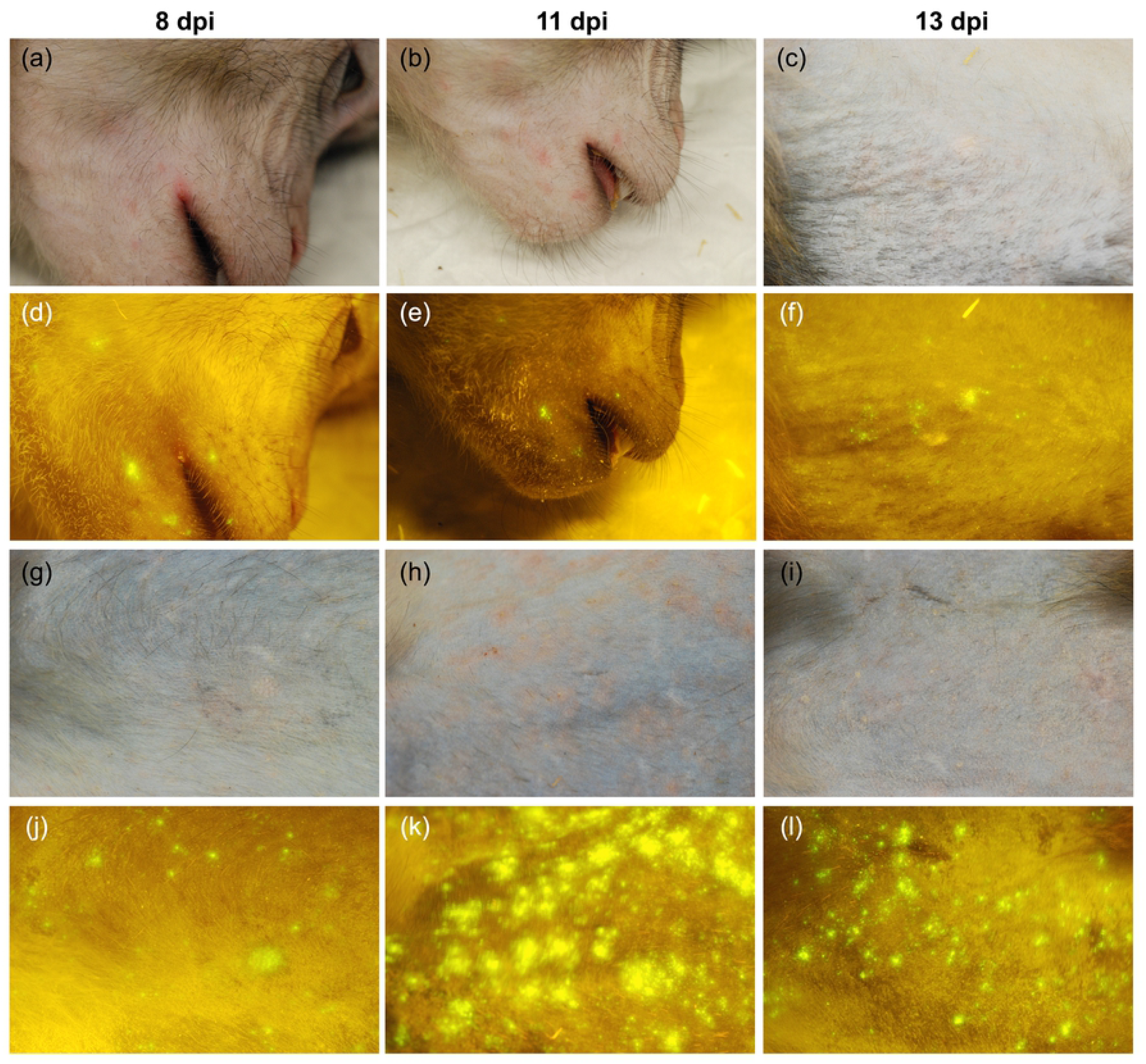
The appearance of MV-infected cells in the skin precedes the appearance of rash. Macroscopic evaluation of MV infection in two cynomolgus macaques: animal #38 (a – f) and animal #37 ((g – l), table S1 in [26]). (a – c; g – i) Normal light: Rash was prominent at 11 dpi. (d – f; j – l) Fluorescence: MV-infected sites (fluorescence) in the skin preceded the rash at 8 dpi and diminished around 13 dpi.

### Infection of dermal immune cells precedes infection of epidermal keratinocytes in experimentally-infected NHPs

We performed immunohistochemistry on formalin-fixed and paraffin-embedded skin samples from these NHPs and showed co-localisation of EGFP and MV nucleoprotein (N) signals in sequential skin sections, which indicated the presence of MV-infected cells. At 9 dpi, the infected cells were predominantly located in the superficial dermis, most especially around the hair follicles and sebaceous glands (Fig 2a – f). Most infection in the epidermis appeared as single-cell infection near dermal papillae and later progressed into multiple-cell infection and syncytia (Fig 2g – i) between 9 and 11 dpi. This epidermal infection later reached the superficial side at 13 dpi, by which time the infection in the dermis already had been cleared. Oedema and spongiosis could be observed at this time point.

**Fig 2.**
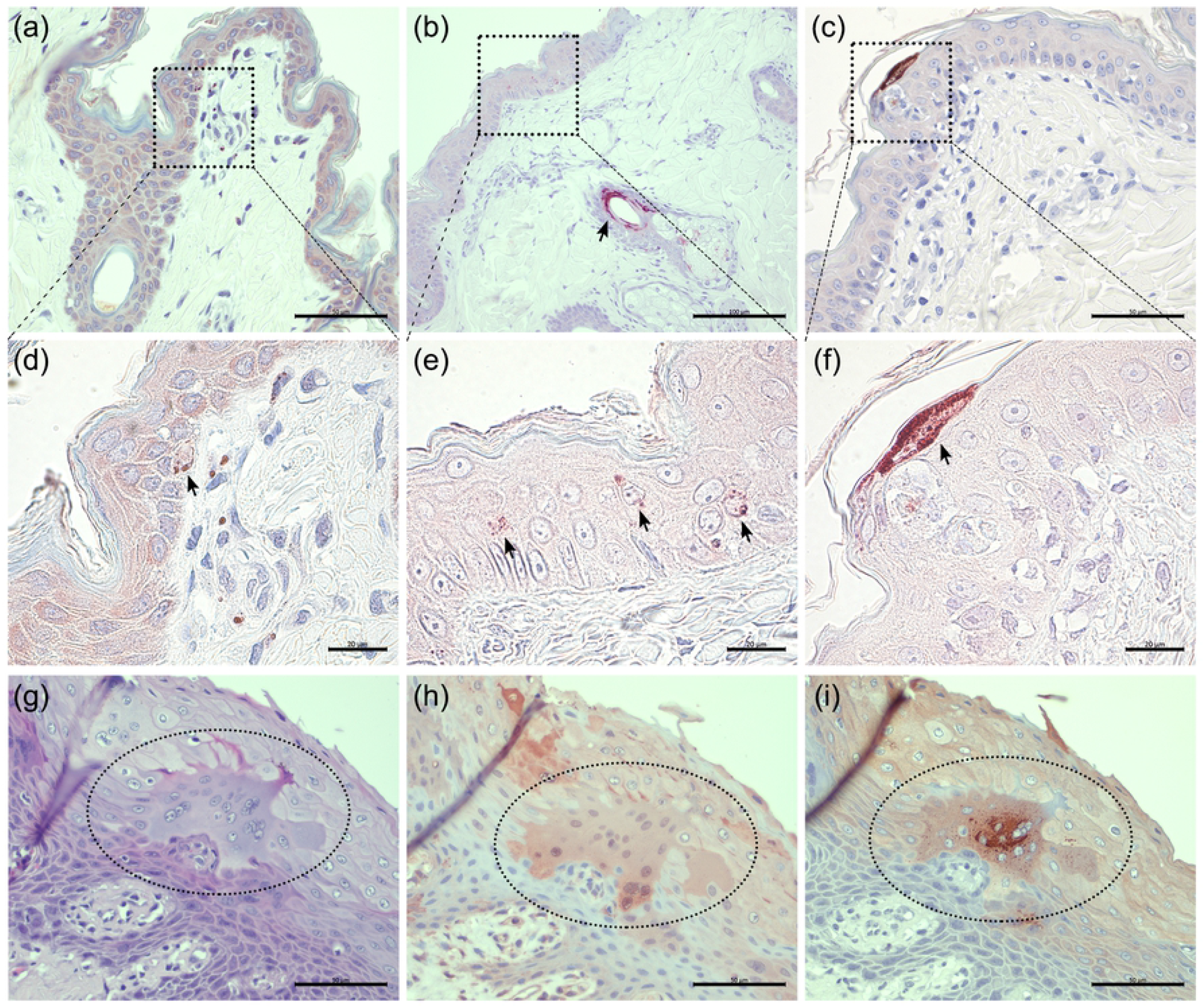
Infection of macaque skin starts in the dermis and spreads to the epidermis. Immunohistochemical staining of MV-infected macaque skin biopsies collected at 9 (a and d), 11 (b and e) and 13 (c and f) dpi. (a and d) Most MV N^+^ cells could be found in the dermal papillae, although a few single infected cells were detected in the basal layer of the epidermis at 9 dpi. (b and e) At 11 dpi, prominent infection was observed near hair follicle and sebaceous gland (arrow). The infection in the epidermis has progressed further in the suprabasal layers (arrows). (c and f) The infection in the dermis was no longer detected at 13 dpi. The infection in the epidermis had reached the most superficial layers. (g – i) A syncytium (ellipse) in the epidermis of macaque skin collected at 9 dpi stained with haematoxylin and eosin (HE), or with green fluorescence protein (GFP) and MV N antibodies, respectively. Scale bars of (a), (c) and (g – i): 50 μm; Scale bar of (b): 100 μm; Scale bars of (d – f): 20 μm.

To identify the phenotype of the infected cells, we performed dual-indirect immunofluorescence (IIF) on these sequential skin sections (Fig 3). CD45^+^ white blood cells were present in the superficial dermis, most especially in or around blood vessels, hair follicles and sebaceous glands and, to a lesser degree, in the epidermis. Some of these CD45^+^ white blood cells were CD3^+^ T cells that were located in the reticular dermis, while some others were S100A8/A9-complex^+^ (Mac387) macrophages that were abundantly present in the superficial dermis, most especially in or around the blood vessels, hair follicles or sebaceous glands. Cytokeratin^+^ cells were restricted to the epidermis and pilosebaceous units. In contrast, CD31^+^ endothelial cells of the blood vessels were restricted to the dermis.

**Fig 3.**
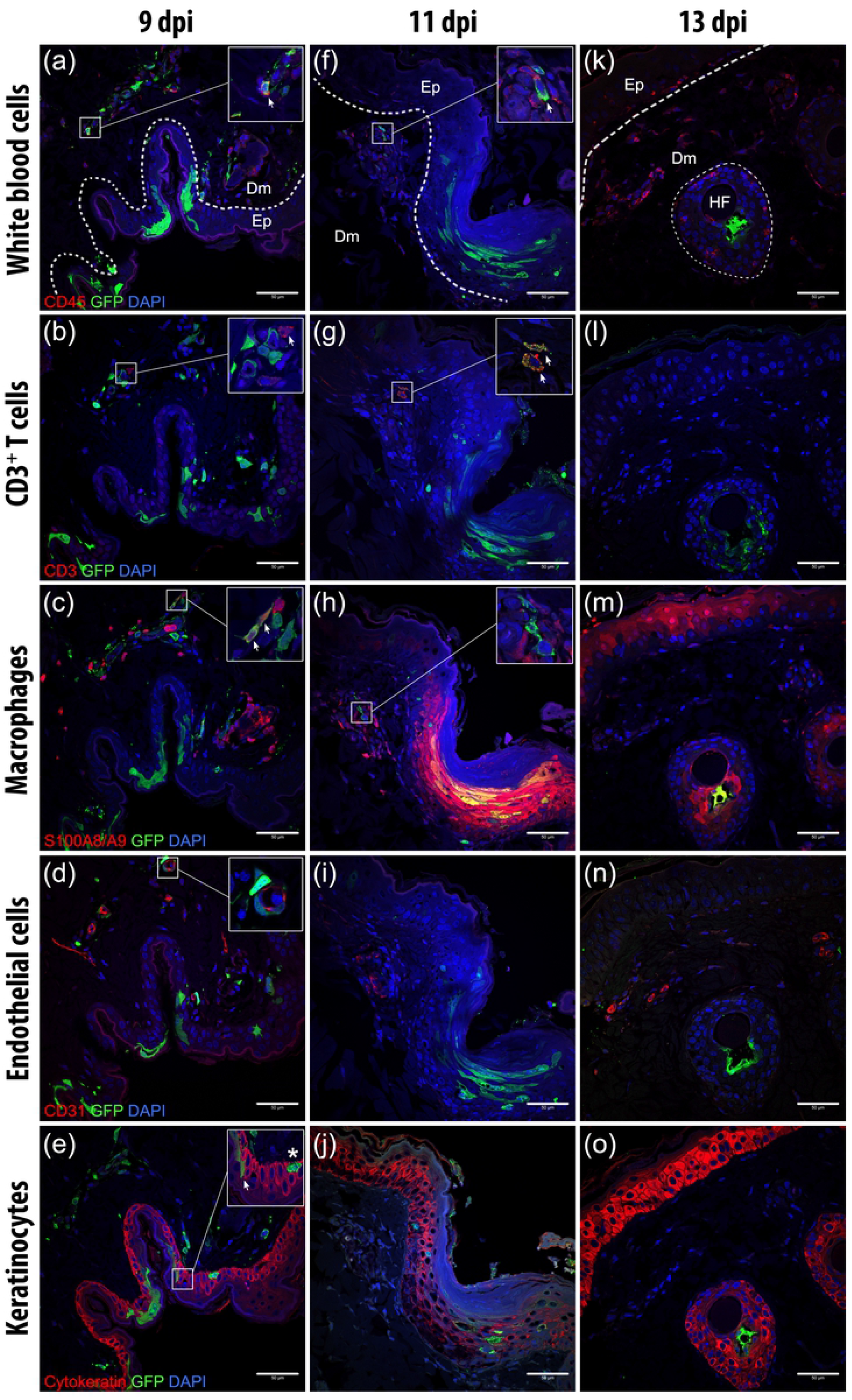
The phenotypes of MV-infected cells in the dermis and epidermis throughout the course of infection. Serial sections of skin (top to bottom) of three macaques (left to right) euthanised at three different time points (indicated above). The sections were double-stained with antibodies to EGFP (green) and several cell-specific markers (red), as indicated on the left of each row. (a) At 9 dpi, MV-infected CD45^+^ white blood cells (inset, arrow) could be detected in the superficial dermis. (b) Some of these MV-infected white blood cells were CD3^+^ T cells, which were present in the dermis, mostly in a more basal area, with speckled GFP signal in their cytoplasm (inset, arrow). (c) MV-infected S100A8/A9 complex^+^ macrophages (inset, arrow) were also found abundantly in the superficial dermis. (d) MV-infected cells in the dermis were often found in or around CD31^+^ blood vessels (inset). (e) In the epidermis, MV-infected cells were mostly keratinocytes (inset, arrow), although MV-infected non-keratinocyte cells (inset, asterisk) were observed in the basal epidermis. (f) At 11 dpi, MV-infected white blood cells (inset, arrow), which were (g) T cells in the dermal papillae (inset, arrow) were in close proximity to (h) uninfected macrophages (inset) and (i) blood vessels. (j) The infection in keratinocytes had progressed apically and laterally. (k – o) MV-infected cells had mostly disappeared from the dermis at 13 dpi. The dermis and epidermis were filled with white blood cells. Dm: Dermis; Ep: Epidermis. Scale bar: 50 μm.

MV-EGFP^+^ cells could be found as early as 9 dpi in the dermal papillae. These cells predominantly belonged to the white blood cell phenotype (Fig 3a), which were mostly T cells or macrophages (Fig 3b – c). Some MV-infected cells were present in or surrounding blood vessels (Fig 3d). In some areas in which the infection was more progressed, the infection spread to the epidermis, even into the most superficial layers, which comprised of differentiated keratinocytes (Fig 3e). The MV-infected white blood cells were still detectable in the dermis on day 11, sometimes in close proximity with uninfected white blood cells, mostly macrophages (Fig 3f – h). These cells clustered close to the dermal papillae, where a lot of blood vessels could be found (Fig 3i). Meanwhile, the infection in the epidermis had progressed laterally and apically (Fig 3j). We also observed keratinocytes at the site of infection expressing S100A8/A9 complex (Fig 3h) as a response to inflammatory stimuli. By 13 dpi, the dermis was almost clear of the infected cells and was filled with white blood cell infiltrates, mostly macrophages (Fig 3k – m). No MV-infected endothelial cells were observed at this time point (Fig 3n). Infection in the epidermis had mostly resolved, although remaining infected follicular keratinocytes could still be detected (Fig 3o).

Focal MV skin infection, in which the progression of infection was different in different sites, was observed as early as 9 dpi. More infected cells were observed in the dermis and epidermis in the sites where the infection had progressed further. At the same time point, but in a different site, where the infection has not progressed, individual MV-infected cells were more often found only in the dermis. This suggested that time was not the only important factor in the pathogenesis of MV skin rash. The location in which MV-infected cells could be found and the interaction with surrounding cells could also play a crucial role in establishing infection in the skin. We observed MV-infected white blood cells in dermal papillae or close to the basal epidermis (Fig 4a). MV-infected T cells, although could often be found near the dermal papillae around 9 and 11 dpi, became scarce at 13 dpi and could only be observed in the reticular dermis. At this late time point, the MV-infected T cells were found surrounded by uninfected T cells (Fig 4b). Close interaction could also be observed among MV-infected cells with HLA-DR^+^ antigen-presenting cells (APCs), for example through a long, EGFP^+^ dendrite (Fig 4c). We observed MV-infected cells surrounded by or in close proximity to endothelial cells (Fig 4d) at 9 dpi at the site where the infection has progressed further. Very rarely, in the same site, we found MV-infected CD31^+^ endothelial cells near other infected cells (Fig 4e). We also observed MV-infected cells in the dermis that were negative for white blood cell, APC, endothelial cell and epithelial cell markers and appeared to be spindle- or dendritic-like cells (Fig 4f).

**Fig 4.**
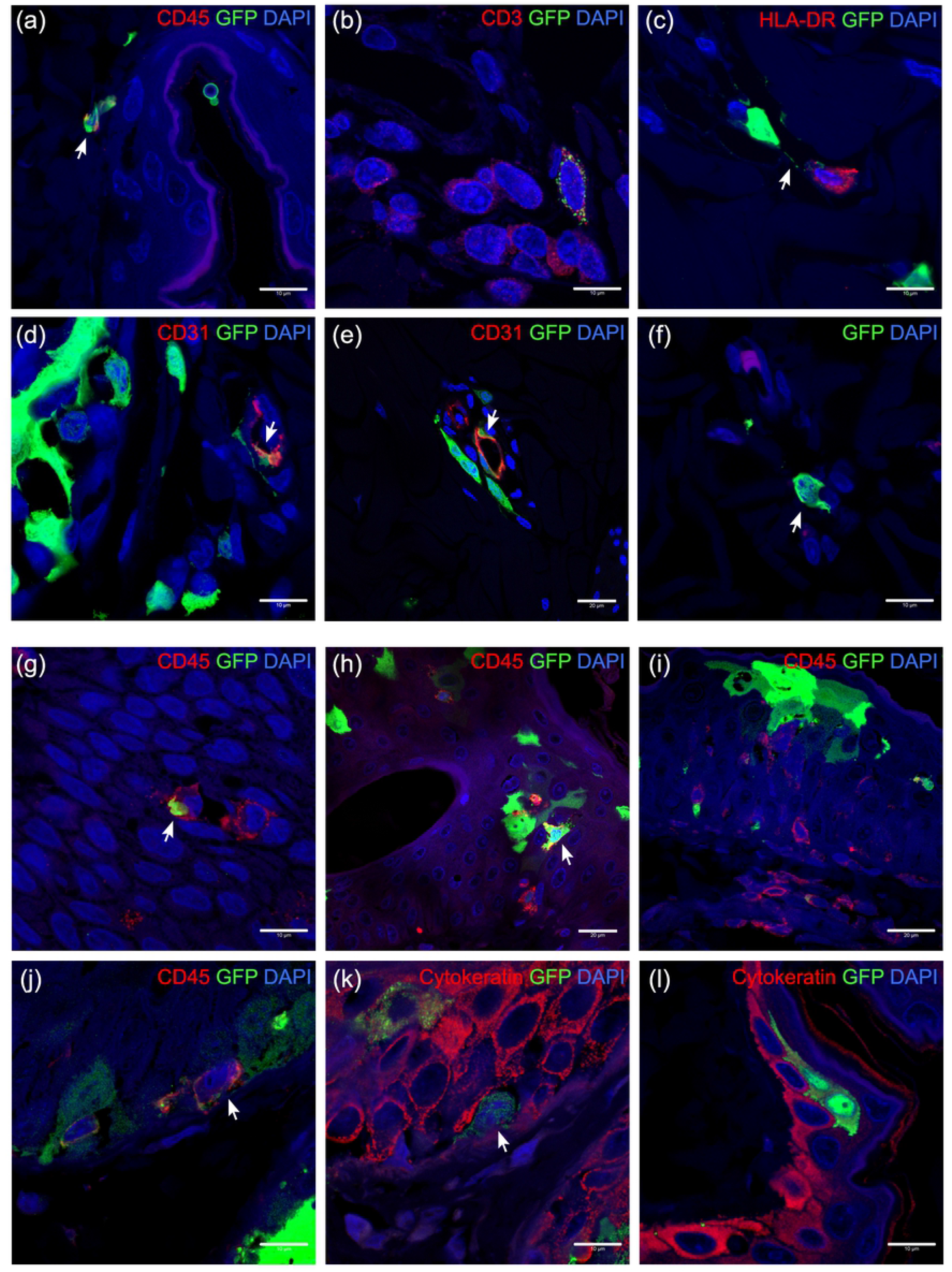
The location of MV-infected cells and the interaction with other cells in their proximity. (a – f) MV-infected cells in the dermis and (g – l) in the epidermis. (a) MV-infected CD45^+^ white blood cells (arrow) in the dermis, especially near the basal layer of the epidermis. (b) MV-infected CD3^+^ T cells, although mostly found in the dermal papillae at 9 and 11 dpi, in reticular dermis at 13 dpi, surrounded by uninfected T cells. (c) MV-infected cell in the dermis interacted with HLA-DR^+^ APC, forming a long EGFP^+^ dendrite (arrow). (d) More often, MV-infected cells (arrow) located around or in blood vessels and, (e) rarely, MV-infected endothelial cells (arrow) could be found together with those cells. (f) Spindle- or dendritic-like MV-infected cells were negative for all tested cell markers. (g – i) In the epidermis, MV-infected white blood cells could be found since 11 dpi, either interacting with other white blood cells (g) or other MV-infected epidermal cells (h). (i) White blood cells appeared to infiltrate the MV-infected cells in the epidermis. (j – k) Serial slides of MV-infected epidermis at 13 dpi. MV-infected white blood cells that were negative for cytokeratin marker could be found in the basal layer of the epidermis, presumably Langerhans cells. These cells were in close proximity with infected keratinocytes (k). (l) MV-infected keratinocytes in the absence of other cells. Scale bars of (a – d), (f), (g) and (j – l): 10 μm; Scale bars of (e) and (h – i): 20 μm.

In the epidermis, a number of MV-infected white blood cells could be observed since 11 dpi, accompanied by infiltration of uninfected white blood cells to the site of infection (Fig 4g – i). These MV-infected white blood cells were negative for keratinocyte, macrophage and T cell markers, thus most likely to be LCs (Fig 4j – k). These cells were found in close proximity to keratinocytes, which were also positive for MV infection (Fig 4k), although keratinocyte infection could still be detected despite the absence of MV-infected white blood cells in the observed two-dimensional plane (Fig 4l).

### *Ex vivo* MV infection of human skin sheets results in higher infection levels in the dermis than in the epidermis

Based on the observations in NHPs, we hypothesised that infection of immune cells in the dermis plays a major role in the pathogenesis of measles skin rash. To test this hypothesis, we *ex vivo* inoculated human full skin or enzymatically-separated epidermal and dermal sheets with rMV based on a wild-type virus strain Khartoum-Sudan (KS) expressing the fluorescent reporter protein Venus from an additional transcription unit in position 3 of the viral genome (rMV^KS^Venus(3)). We monitored the infection up to seven days. We observed that Venus^+^ cells could be detected by inverted laser scanning microscopy as early as 2 dpi, with higher infection levels in the dermis than the epidermis (Fig 5a). The phenotype and the percentage of emigrant MV-infected cells from the skin sheet cultures were determined by flow cytometry. These cells turned out to be CD4^+^ T cells, APCs and non-lymphocytes. In accordance to the observation on Venus^+^ cells under the inverted laser scanning microscopy, the percentage of MV-infected cells was higher in the dermis than in the epidermis (Fig 5b). This finding suggested that the dermis is an important site where MV infection is established.

**Fig 5.**
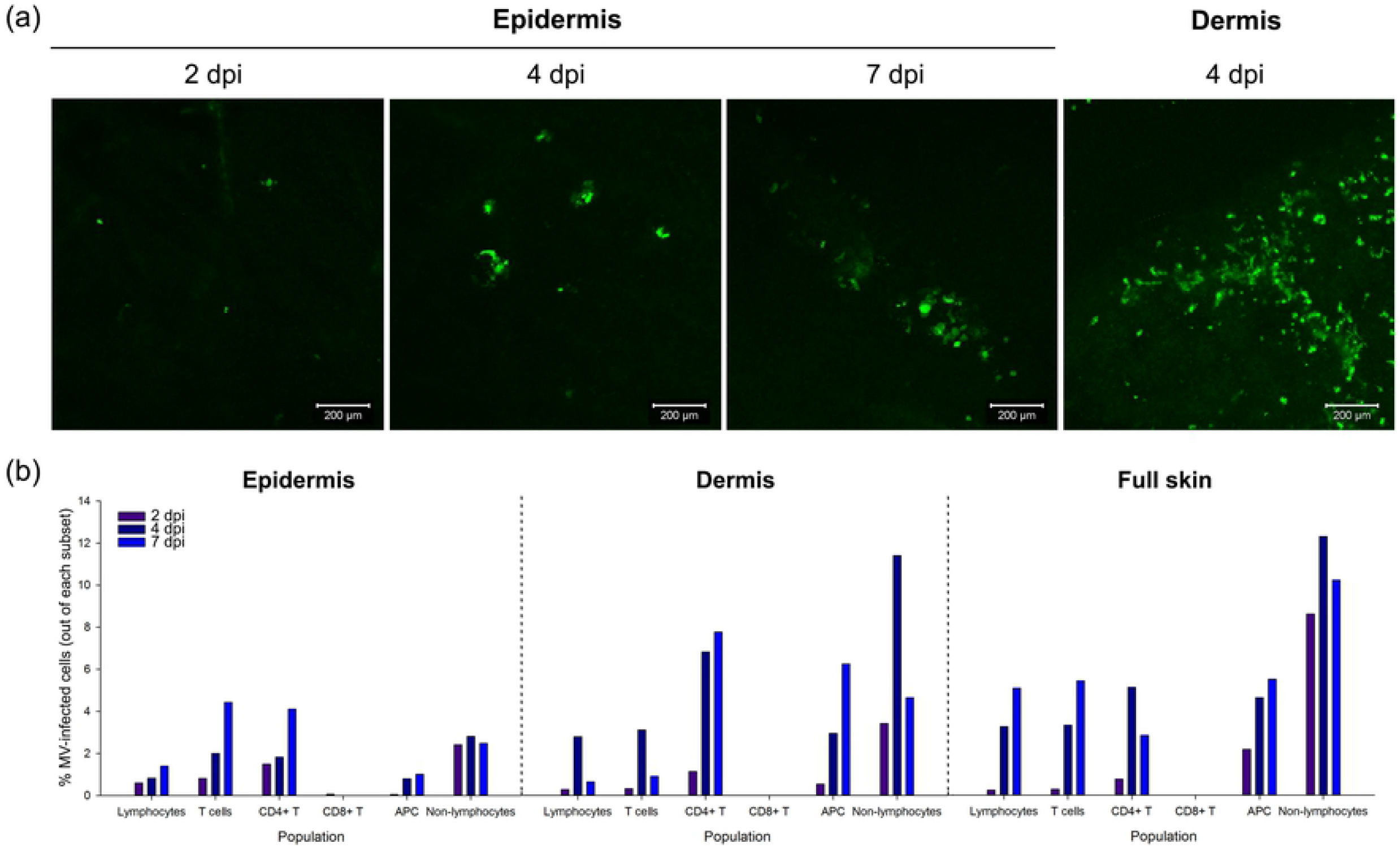
*Ex vivo* MV infection of human epidermis and dermis sheets. (a) MV^+^ cells (green) were detectable as early as 2 dpi in the epidermis and were present in higher numbers in the dermis than in the epidermis. (b) Percentages of infected cells in each cell subset of lymphocyte and non-lymphocyte populations isolated from epidermis, dermis and full skin at 2, 4 and 7 dpi. APC: antigen-presenting cell. Scale bars: 200 μm.

Previous studies showed that mature LCs are susceptible to MV infection and Langerin can act as an attachment receptor, but not entry receptor, for the virus [28]. To determine whether LCs play a role in MV epidermal infection as initial target cells, we performed dual-IIF stainings on human epidermal sheets from healthy donors infected with rMV^KS^Venus(3). We found that despite the abundant presence of LCs in the epidermal sheets, none of these were Venus-positive (S1). This finding suggests that LCs do not play a major role in the pathogenesis of measles skin rash.

### Human primary keratinocytes are susceptible to *in vitro* MV infection in a nectin-4-dependent manner

To investigate the susceptibility and the permissiveness of keratinocytes, we inoculated human primary human keratinocytes derived from two healthy donors or from a patient affected by ectodermal dysplasia-syndactyly syndrome (EDSS1, OMIM 613573), an autosomal recessive disorder caused by mutations in the nectin-4 encoding gene *PVRL4* [25]. In this patient, EDSS is secondary to compound heterozygous mutations c.554C>T and c.906delT in the *PVRL4* gene (family B in [25]). While the first is a missense mutation (p.Thr185Met) affecting a highly conserved residue of nectin-4 located in its second extracellular immunoglobulin-like domain, the second variant leads to a prematurely truncated protein (p.Pro304HisfsX2). Previous study has shown that in cultured epidermal keratinocytes from this EDSS patient, nectin-4 mRNA displayed nearly 50% reduced expression in line with nonsense mRNA decay of the c.906delT allele, suggesting that all the residual protein expressed on the keratinocyte surface (about 10%) is represented by the p.Thr185Met mutant [25]. In agreement with previously published data, nectin-4 expression on the cell surface was highest in differentiated healthy donors’ keratinocytes, as demonstrated by flow cytometry. However, differentiation did not result in increase of nectin-4 expression in the EDSS patient’s keratinocytes (S2) [25]. To determine whether the proliferating and differentiated keratinocytes were susceptible to MV infection, we inoculated them with two MV strains expressing fluorescent reporter proteins (rMV^IC323^EGFP(1) or rMV^KS^Venus(3)) [29] or a strain engineered to be unable to recognise nectin-4 (the ‘nectin-4-blind (N4B)’ strain rMV^KS-N4b^EGFP(3)) [10] at a multiplicity of infection (MOI) of 1. After 48 hours, we observed higher frequencies of fluorescent cells in differentiated than in the proliferating cells (Fig 6). Infection of keratinocytes with the nectin-4-blind MV resulted in low numbers of single infected cells. Few MV infected cells were detected in the EDSS proliferating and differentiated cultures.

**Fig 6.**
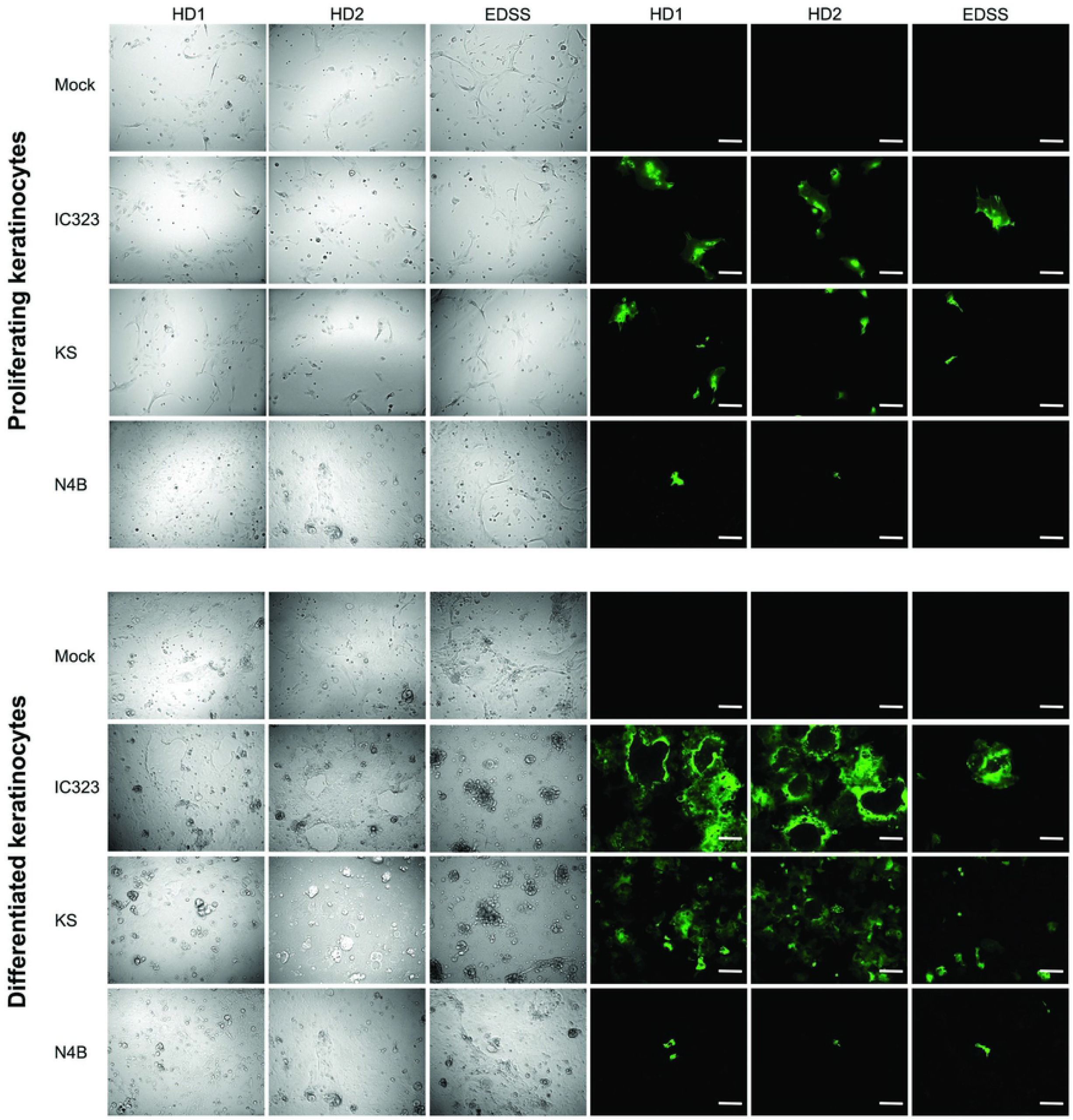
The susceptibility of proliferating and differentiated keratinocytes to *in vitro* MV infection. A higher number of infected keratinocytes (green) was detected in differentiated than in proliferating cultures, regardless of MV strain. All experiments were done in duplicate. NCI-H358: human broncho-alveolar carcinoma cell line; BLCL: EBV-transformed B-lymphoblastoma cell line. HD1 or HD2: primary keratinocyte culture from healthy donor 1 or 2; EDSS: primary keratinocyte culture from EDSS patient; KS: rMV^KS^Venus(3); IC323: rMV^IC323^EGFP(1); N4B: rMV^KS-N4b^EGFP(3). Scale bars: 200 μm.

To assess whether the infected keratinocytes also produced cell-free virus and were thus capable of spreading the infection, the supernatant of the MV-infected keratinocytes was collected and the titre of cell-free virus in the supernatant was assessed [30]. Cell-free MV was detectable in the culture supernatant of the infected proliferating and differentiated keratinocytes. In healthy donors, virus titres in supernatants of differentiated keratinocytes were higher than those in supernatant of proliferating keratinocytes. Interestingly, virus titres of the EDSS patient’s proliferating keratinocytes were comparable to those of the healthy donors (Fig 7a). The difference in virus titres increased as the cells differentiated, since the titre was higher in healthy donors than in the patient.

**Fig 7.**
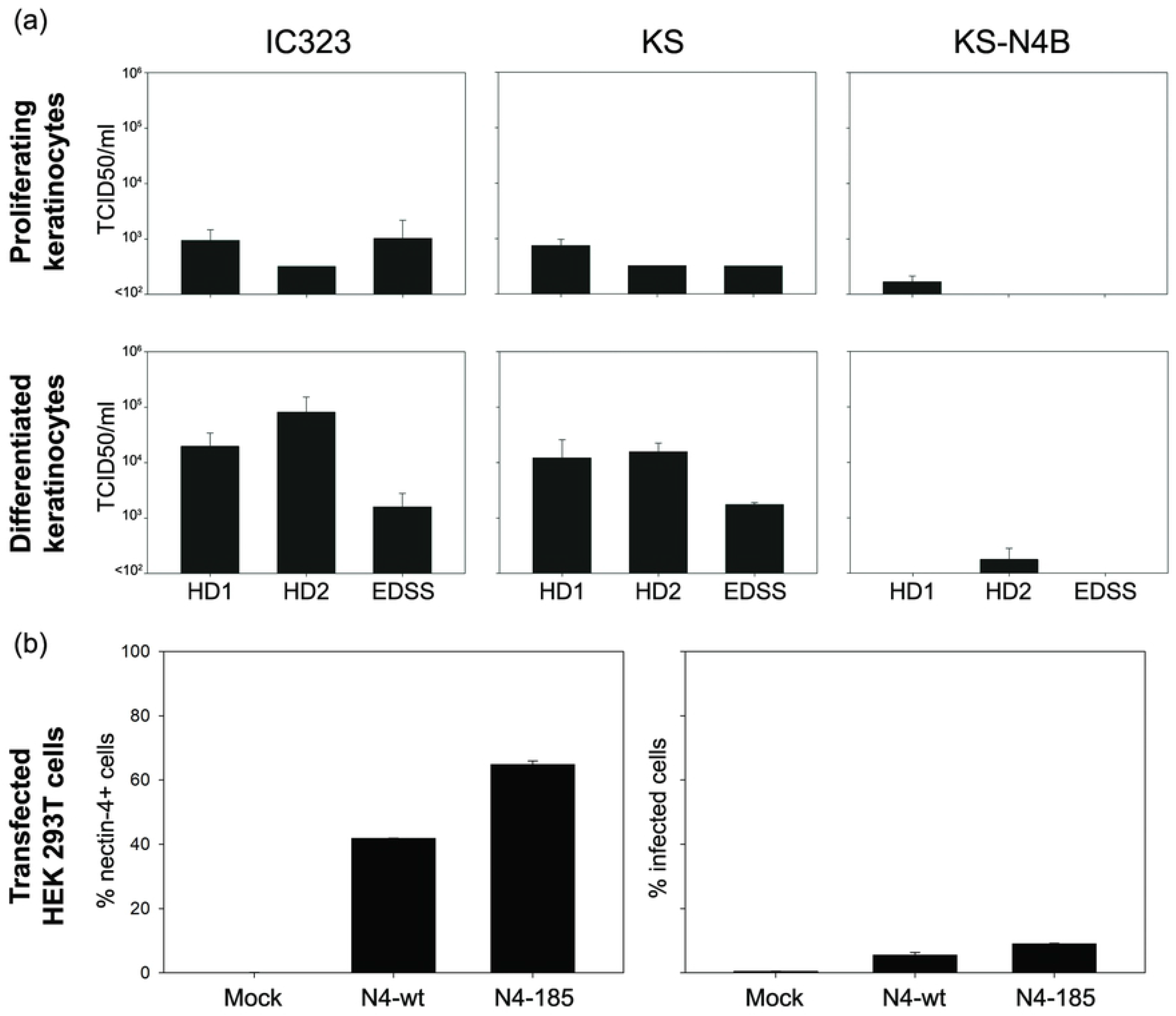
Production of cell-free virus by MV-infected proliferating and differentiated keratinocytes and susceptibility of HEK 293T cells transiently expressing wild-type or mutant nectin-4 to MV infection. (a) MV-infected proliferating and differentiated keratinocytes of an EDSS patient produced a comparable amount of cell-free virus to that of the healthy donors. (b) Expression of nectin-4 on HEK 293T cells was measured prior to infection (left graph). Venus fluorescence was detected in these cells after 24 h of infection (right graph). Mock cells were taken as control for transfection and infection. HEK 293T cells were transfected with plasmid containing wild-type or mutant p.Thr185Met insert. All experiments were done in duplicate. HD1 or HD2: primary keratinocyte culture from healthy donor 1 or 2; EDSS: primary keratinocyte culture from EDSS patient; KS: rMV^KS^Venus(3); IC323: rMV^IC323^EGFP(1); KS-N4B: rMV^KS-N4b^EGFP(3); N4-wt: wild-type nectin-4; N4-185: nectin-4-T185M.

Low expression of mutant nectin-4 on the surface of EDSS patient’s keratinocytes could apparently still facilitate MV infection, leading to the production of new viral particles by infected keratinocytes, suggesting that the p.Thr185Met mutation in the second immunoglobulin-like domain does not affect the virus binding. To validate this observation, we transiently transfected human embryonic kidney (HEK) 293T cells with a plasmid containing either wild-type or two mutant nectin-4 (p.Thr185Met) insert. The cells were then inoculated directly with rMV^KS^Venus(3) at an MOI of 3. Infection efficiency was measured by flow cytometry 24 hours post-infection. Expression levels of both wild-type and mutant nectin-4 was comparable and both nectin-4-transfected cells, but not the mock-transfected cells, proved susceptible to MV infection (Fig 7b), which confirmed that the defective nectin-4 (p.Thr185Met) could still function as a cellular receptor for MV.

Based on our findings, combined with previously published observations, we postulate a model that describes the progression of MV skin infection and the development of measles rash (Fig 8). The model takes viral tropism, location, interaction and motility of the susceptible cells, as well as the virus-specific immune responses into account. MV-infected cells enter the superficial dermis through the blood vessels and spread the infection to the tissue-resident dermal T cells, APCs and spindle- or dendritic-like cells around 7 dpi. The infection progresses several days later to the adjacent epidermal areas, where the infection is transmitted to the basal keratinocytes. As basal keratinocytes differentiate vertically to the suprabasal layers and their nectin-4 expression increases, the virus spreads laterally and the infected keratinocytes subsequently form syncytia. Infection of dermal endothelial cells was very rare, but not completely absent. We speculate that the infection is subsequently cleared around 13 dpi by infiltrating MV-specific T cells, which first migrate into the dermis and later into the epidermis.

**Fig 8.**
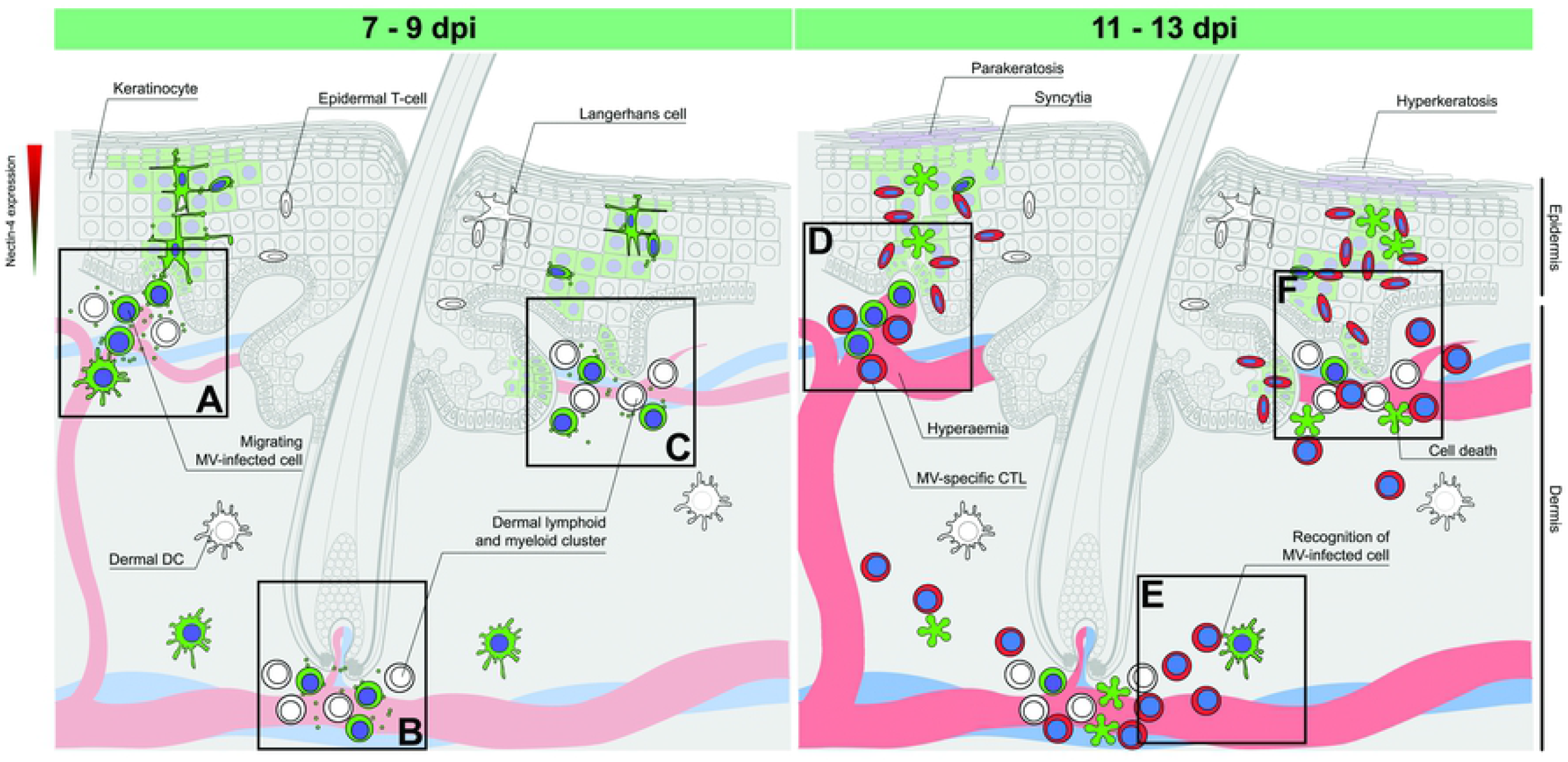
Model for pathogenesis of measles skin rash. During viremia, MV-infected T cells and macrophages migrate to the dermis via the capillaries and interact with (a) tissue-resident lymphoid and myeloid cells and epidermal LCs residing near the basal lamina. This interaction leads to the infection of surrounding CD150^+^ tissue-resident immune cells and nectin-4^+^epithelial cells. Alternatively, MV-infected T cells and macrophages migrate in close proximity to: (b) the hair follicle or (c) the sebaceous gland via the capillary, where they infect an aggregate of tissue-resident T cells and macrophages, and further spread the infection to nearby keratinocytes and LCs. Infection of basal keratinocytes leads to lateral and apical spread of the virus to the superficial layers of the epidermis. Several days later, (d) hyperaemic responses allow the recruitment of MV-specific CD8^+^ cytotoxic T cells and macrophages, resulting in (e) recognition and (f) clearance of the infected cells. Hyperaemia and subsequent oedema are the histological correlates of maculopapular erythematous measles rash.

## Discussion

The pathogenesis of MV skin infection and subsequent skin rash is not well understood. Here, we aimed to identify the cell types involved in MV infection of skin, and the kinetics of viral dissemination in relation to onset of rash. Based on observations from experimentally infected NHPs, *ex vivo* infected human skin explants and *in vitro* infected primary keratinocytes, we initially observed MV-infected perivascular, perifollicular and periadnexal dermal immune cells, followed by dissemination to epidermal keratinocytes. Skin infection preceded onset of rash by 3 to 4 days.

The dermis contains several potential target cells for MV infection. Due to the blood vessels and capillaries that run through it, the dermis is filled with CD150^+^ lymphoid and myeloid cells that traffic through or reside in the tissue. CD4^+^ and CD8^+^ T cells localise and move differently in the skin [31]. Slow-moving CD8^+^ resident memory T cells (T_RM_) reside in the epidermis and hair follicles, while highly motile CD4^+^ effector memory T cells (T_EM_) migrate into the dermis and recirculate systemically [32]. We detected MV-infected T cells in the dermis from 9 dpi onward, but never in the epidermis at that time point. Previous studies have shown that CD4^+^ TEM cells are highly susceptible to MV infection [26, 33]. Interaction of MV-infected T cells with skin-resident APCs may result in further cutaneous spread. T cells have been described in human skin to cluster with APCs around appendages, such as hair follicles [34–36]. We did not observe such T cell clusters in NHPs, most likely due to T cell depletion that occurs systemically and peaks around 9 dpi [26]. Whether this depletion leads to the loss of pre-existing skin-resident memory T cells remains to be studied. Additionally, we and others have observed MV-infected T cells and APCs around hair follicles and sebaceous glands [15, 37], which are surrounded by nectin-4^+^ epithelial cells [25]. The close proximity of these infected cells to the basal keratinocytes may lead to the spread of MV infection from the dermis to epidermis.

The epidermis consists predominantly of keratinocytes, which express nectin-4 and are susceptible to MV infection [24]. We were not able to determine the expression of nectin-4 in the NHP epidermis due to the lack of cross-reactive antibodies. However, in accordance with the previous study, we show primary human keratinocytes express nectin-4 and its expression is upregulated upon differentiation [25]. We show here that nectin-4 expression plays a role in the susceptibility of keratinocytes to *in vitro* MV infection: higher expression of nectin-4 resulted in higher susceptibility. This led to the question whether the keratinocytes of an EDSS patient, which have strongly reduced expression of nectin-4, were susceptible to MV infection. We found that, despite its very low expression, the nectin-4 mutant could still facilitate MV infection *in vitro*. Residual nectin-4 activity in this EDSS patient therefore still allows binding of MV haemagglutinin protein and hence facilitates viral entry. Notably, the missense mutation p.Thr185Met is located in the second immunoglobulin-like domain, while MV-binding interfaces of nectin-4 are located in three loops of its most external domain.

Another cell type of interest in the epidermis is the LC, a subset of DCs. Although we could not observe MV-infected LCs in our human skin *ex vivo* model, LCs are known to be susceptible to MV infection [38–40]. The activation status of the cells also determines their susceptibility, since immature LCs are not susceptible to MV infection, while mature ones are [28]. This offers an explanation to why the LCs were not susceptible to MV infection in our *ex vivo* model: the cells might still have been in their immature state. We were also not able to identify LCs in cynomolgus macaque skin tissues due to the unavailability of cross-reactive antibodies. The susceptibility of LCs to MV infection *in vivo* and their role in the pathogenesis of measles skin rash remain to be determined. Additionally, LCs express Langerin that can act as an attachment, but not entry, receptor to MV [28] and thus can indirectly introduce MV infection to the epidermal keratinocytes by acting as an attachment hub for the virus from the dermis. Although we were able to clearly identify APCs and T cells in the dermis, we were not able to find HLA-DR^+^ or CD3^+^ cells in the epidermis. The involvement of these cells in the pathogenesis of MV infection in the epidermis remains to be elicited.

DCs and macrophages occupy the dermis as professional APCs and phagocytes, respectively. Macrophages are present in high numbers and are associated with blood or lymphatic vessels, while dermal DCs have been found to form clusters with T cells, suggesting the presence of an inducible structure of macrophages, DCs and T cells that may function as a skin-associated lymphoid tissue [41, 42]. In the respiratory tract, DCs and macrophages act as Trojan horses during MV infection by spreading the virus to the lymphocytes in draining lymph nodes [7, 43–46]. Migrating or patrolling MV-infected DCs and macrophages may play the same role in the skin as they do in the respiratory tract. However, these cells may also play a crucial role as innate immune cells that inhibit infection. Close communication of MV-infected DCs and macrophages with T cells can lead to activation of MV-specific immune responses and subsequently to the development of rash. The role of these immune responses in the development of rash has been highlighted in immunocompromised patients with MV infection that do not develop skin rash [14].

Blood vessels and capillaries run through the dermis. The capillaries penetrate into the dermal papillae, from where the distance to the epidermis is minimal, and the distribution of the capillary loops differs according to the type of the skin. The capillary bed consists of an arteriole, which gives rise to metarterioles and subsequently hundreds of capillaries. The capillaries provide the dermis and epidermis with nutrition and oxygen, and connect to venous capillarioles and further to a venule. Inflammation due to infection may cause prolonged vasodilatation and increased capillary permeability. This hyperaemic reaction allows the release of chemokines by skin-resident cells, such as memory immune cells and keratinocytes, that leads to the infiltration of various immune cells, such as macrophages and lymphocytes. The vasodilatation also causes erythema and oedema [47]. Given that measles rash is described as maculopapular (*i.e*. small with raised bumps) and erythematous (*i.e*. red), and oedema can be observed in MV-infected skin [15], we speculate that hyperaemia is responsible for the appearance of the erythematous maculopapular rash. Although theoretically it is possible to investigate the presence of hyperaemia in our *in vivo* model by showing an increased number of erythrocytes in the cutaneous blood vessels, we could not perform the calculation fairly, since the animals were sacrificed by exsanguination.

MV infection in the skin gives a unique appearance of rash compared to other viral exanthemata. Rubella rash, for example, has been described as macroscopically similar to measles rash, since it gives a pink-reddish “rubelliform” maculopapular rash. However, in rubella, viral infection takes place deeper in the dermis, in contrast to measles skin infection that occurs more superficially in the dermis. Infection of the keratinocytes, which is typical for measles rash, does not occur during rubella virus infection [48]. In contrast, varicella zoster virus (VZV), as a representative of the *Herpesviridae* family member, has similar target cells in the dermis and epidermis as MV, but displays a different type of rash. VZV infects perivascular macrophages and DCs as well as keratinocytes, but the infection leads to the appearance of spots that turn into itchy blisters [49]. Arboviral exanthemata, on the other hand, have a different route of infection, but often present overlapping outcomes in the skin. Dengue virus is introduced into the body through a mosquito bite and injected into the bloodstream, with spillover to the epidermis and the dermis. This spillover causes infection of LCs and keratinocytes. Dengue virus spreads systemically through the infection of monocytes and macrophages. The virus also causes vascular leakage through infection of endothelial cells, leading to the appearance of minor haemorrhagic lesions [50]. Although petechial rash is one of the clinical manifestations of dengue virus infection, morbilliform rash is also often described during classical dengue fever [51]. Altogether, these findings, including ours, strongly suggest that the appearance of rash, especially during measles, is closely linked to the viral tropism, the availability and location of susceptible target cells and the subsequent immune responses to clear the infection.

MV infects the respiratory epithelial cells and is shed apically into the mucus lining the lumen of the upper and lower respiratory tract, which is void of CD150 or nectin-4. The virus is thus transported to the throat by the mucociliary escalator and expelled into the air by coughing [52]. The role of MV skin infection in viral transmission is still a subject of speculation. The outer layer of the epidermis consists of dead keratinised cells. Whether these dead cells allow the attachment of MV and hence the release of dead-cell-associated virus particles into the air remains to be investigated [15].

In conclusion, our study offers a new explanation to the pathogenesis of measles skin rash: MV-infected lymphocytes and myeloid cells enter the dermis, where the infection spreads to the susceptible cells in the vicinity of dermal papillae, hair follicles, sebaceous glands and blood vessels in the superficial dermis. The infection spreads laterally and apically to the epidermis in a nectin-4-dependent manner. The infection is cleared several days later by infiltrating MV-specific T cells, accompanied by the appearance of oedema and hyperaemia that give the appearance of an erythematous morbilliform rash.

## Materials and Methods

### Ethical statement

All NHP samples were derived from previously published studies, and no new experimental infections were performed [26]. Studies involving the use of primary keratinocytes were approved by the local ethics committee, and written informed consent was obtained from both the EDSS1 patient and the healthy volunteers [25]. Studies using human skin tissue were performed in accordance with the Amsterdam University Medical Centres (AUMC) institutional guidelines with approval of the Medical Ethics Review Committee of the AUMC, location Academic Medical Centre, Amsterdam, the Netherlands, reference number: W15_089 # 15.0103. All samples were handled anonymously.

### Cells

Culture of normal and EDSS primary human keratinocytes was carried out as previously described [25]. Keratinocytes were cultured till sub-confluence in serum-free Keratinocyte Growth Medium (KGM, Invitrogen) containing 0.15 mM Ca^2+^ (proliferating keratinocytes), and then induced to differentiate by culturing for further 3 days in a 3:1 mixture of DMEM and Ham’s F12 media (Invitrogen, Palo Alto, CA) containing FCS (10%), insulin (5 μg/ml), transferrin (5 μg/ml), adenine (0.18 mM), hydrocortisone (0.4 μg/ml), cholera toxin (0.1 nM), triiodothyronine (2 nM), EGF (10 ng/ml), glutamine (4 mM), and penicillin-streptomycin (50 IU/ml) (differentiated keratinocytes). Epstein-Barr virus- (EBV-) transformed B-lymphoblastoma cell line (BLCL) and human broncho-alveolar carcinoma (NCI-H358) cell lines were grown in RPMI-1640 medium supplemented with 10% of fetal bovine serum (FBS), 100 IU of penicillin/ml, 100 μg of streptomycin/ml and 2 mM glutamine (R10F medium). Vero cells expressing human CD150 (Vero-CD150) were grown in Dulbecco’s modified Eagle medium (DMEM) supplemented with 10% of FBS, 100 IU of penicillin/ml, 100 μg of streptomycin/ml and 2 mM glutamine (D10F medium) [53]. Human embryonic kidney (HEK) 293T cells were grown in D10F medium supplemented with 1 mM sodium pyruvate and non-essentials amino acids. All cells were cultured in a humidified incubator at 37° C with 5% of CO2.

### Tissues

Residual skin materials were obtained from six different adult human donors undergoing correctional surgery and stored at 4° C overnight. The skin was shaved using a dermatome (0.3 mm, Zimmer Biomet, UK). For the preparation of full skin sheets, which consist of dermis and epidermis, the shaved skin was cut into circular sheets (diameter approximately 1 cm) using a skin biopsy punch and cultured in IMDM supplemented with 10% of FCS, 100 IU of penicillin/ml, 100 μg of streptomycin/ml (Invitrogen), 2 mM glutamine and 20 μg/ml gentamicine (Centrafarm, Netherlands) (I10F medium), with the epidermis facing upward. The full skin pieces were stored in a 24-well plate in I10F medium. For the preparation of epidermal sheets, shaved skin was incubated in I10F medium in the presence of 1 U/ml of dispase (Roche Diagnostics) for 1 h at 37° C or 0.5 U/ml overnight at 4° C. The epidermis was separated from dermis using a pair of forceps and cut into circular sheets using a skin biopsy punch. The epidermal or dermal sheets were stored in a 24-well plate in I10F medium, with the keratin layer of the epidermis facing upward.

### Viruses

All recombinant MV strains used in this study were described previously: recombinant MV strain Khartoum-Sudan (KS) expressing the fluorescent protein Venus from an additional transcription unit in position 1 or 3 (rMV^KS^Venus(1) or (3)) [54] and strain IC323 expressing the fluorescent protein EGFP in position 1 (rMV^IC323^EGFP(1)) of the viral genome [29] were based on wild type viruses. An rMV^KS^ expressing EGFP in position 3 of the viral genome engineered to be unable to recognise nectin-4 (referred to as the ‘nectin-4-blind (N4b)’ rMV^KS-N4b^EGFP(3)) was also included in this study [55]. Virus titres were determined by endpoint titration on Vero-CD150 cells, and were expressed as 50% tissue culture infectious dose (TCID50) per ml calculated as described by Reed and Muench [30].

### Sequencing

Partial sequences of plasmid 3XFLAG containing wild-type or mutant nectin-4 (p. Thr185Met) insert were obtained with Applied Biosystems 3130xl Genetic Analyser, according to the instructions provided by the manufacturer. The primers used in this assay were: 5’-CCT-GCC-CTC-ACT-GAA-TCC-TG-3’ and 5’-ACA-CCC-ACC-ACC-ACC-ACC-GA-3’. The obtained sequences were analysed with BioEdit software.

### Transfection

De Wit *et al*. have previously described transfection of HEK 293T cells [56]. In brief, prior to transfection, HEK 293T cells (3 × 10^6^) were seeded into gelatinised 10-cm^2^ culture dishes. The cells were transiently transfected overnight in the present of CaCl2 with 3XFLAG plasmid (30 μg) containing wild-type or mutant nectin-4 insert. After 6 h of transfection, the cells were treated with trypsin and seeded into gelatinised 24-well plates. After 36 h of transfection, the cells were treated with trypsin and the nectin-4 expression levels were assessed with flow cytometry.

### *In vitro* MV infection

Adherent primary keratinocytes were either inoculated directly or were treated with trypsin-EDTA (0.05%) and inoculated in suspension with the three different rMV strains at an MOI of 1. After 2 h, the suspension cells were washed to remove unbound virus and seeded onto 24-well plates in K medium. After 48 h of infection, the cells were observed under an inverted-laser scanning LSM-700 microscope (Zeiss) and the infection percentages were assessed by flow cytometry. Transiently transfected HEK 293T cells were inoculated directly with rMV^KS^Venus(3) at an MOI of 3. After 24 h of infection, the infection percentages were measured by flow cytometry.

### Ex vivo MV infection

Full skin pieces, dermal or epidermal sheets were inoculated with cell-free rMV^KS^Venus(3). Briefly, 200 μl of pure virus stock (3.7 × 10^6^ TCID_50_/ml) was added to each well of a 24-well plate, and the skin sheets were added on top of the liquid with the epidermis facing upwards. While full skin and epidermal sheets remained afloat, dermal sheets tended to sink and both apical and basolateral surfaces were exposed to virus. After 2 h at 37°C, I10F medium was added to the wells. The progression of infection was observed at 2, 4 and 7 dpi under the inverted laser scanning microscope. Full skin pieces, dermal or epidermal sheets were fixed in 4% paraformaldehyde (PFA) for at least 24 h and subsequently stored in PBS or as formalin-fixed paraffin-embedded tissue blocks.

### Measurement of MV production by infected keratinocytes

Supernatant of MV-infected keratinocytes was titrated into 96-well plates containing Vero-CD150 cells (1 × 10^4^ cells/well). The titre of the virus was expressed as TCID_50_/ml and calculated as described above.

### Flow cytometry

Flow cytometry was performed using a BD FACSCanto II. Primary keratinocytes or HEK 293T cells were labelled with nectin-4^PE^ antibody (clone 337516; R&D Systems) to assess the expression of nectin-4. Isotype control (Isotype^PE^, clone 27-35, BD Biosciences) antibody was included to assess the level of background staining. NCI-H358 cells and BLCL were included as positive and negative controls for nectin-4 expression, respectively. All cells were fixed with 2% of PFA prior to measurement of the percentage of cells expressing the virus-encoded fluorescent protein. Mock-infected cells were included as infection control. Supernatants from full skin pieces (n = 3 donors), dermal (n = 1 donor) or epidermal sheets (n = 3 donors) were isolated at 2, 4 and 7 dpi and emigrant cells were isolated after undergoing centrifugation. Antibodies used in this experiment is listed in Table 1. Data was acquired with BD FACSDiva software and analysed with FlowJo software.

**Table 1.**
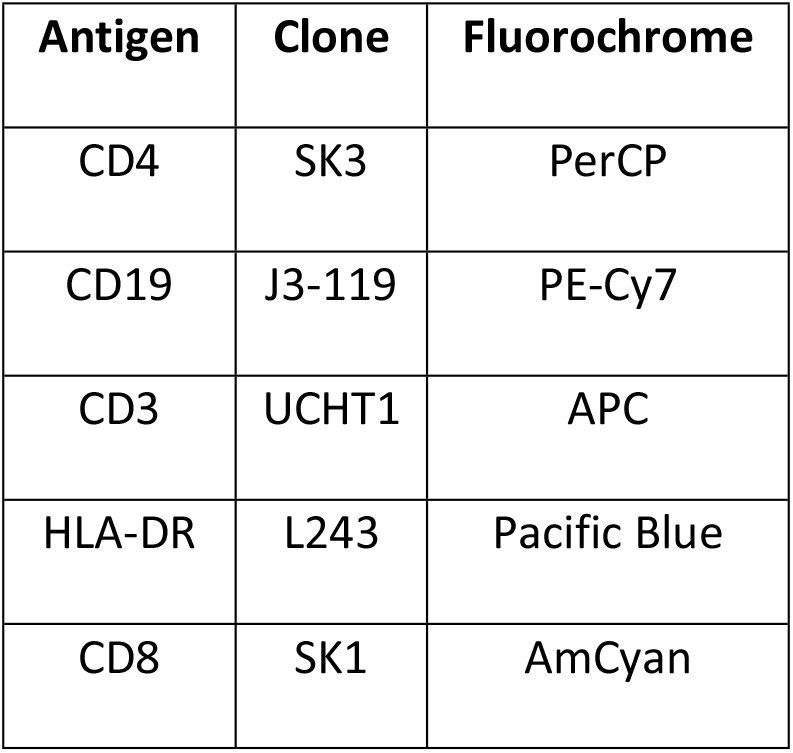
List of antibodies used in flow cytometry to identify the phenotypes of cells in full skin, dermal and epidermal sheets.

### *In situ* analyses

Immunohistochemistry was performed using monoclonal antibodies directed to MV N protein (clone 83KKII, Chemicon [57]) or rabbit polyclonal antibody directed to GFP (Invitrogen). Goat anti-mouse IgG1 or goat anti-rabbit antibody conjugated with biotin was included as secondary antibody. Streptavidin-horseradish peroxidase was added for signal detection. Dual-IF assays were performed using mouse monoclonal antibodies directed to CD45 (clone 2B11+PD7/26; DAKO), CD3 (clone F7.2.38; DAKO), CD31 (clone JC70A; DAKO), cytokeratin (clone AE1/AE3; DAKO), S100A8/A9 complex (clone MAC387; Abcam), or HLA-DR (clone L243; BioLegend) in combination with rabbit polyclonal antibody directed to GFP. Goat anti-rabbit-IgG-Alexa Fluor (AF)488 (Invitrogen) and goat anti-mouse IgG-AF594 (Invitrogen) were included as secondary antibodies. Formalin-fixed, paraffin-embedded tissues were sectioned at 3 μm, deparaffinised and rehydrated prior to antigen retrieval. Antigen retrieval for MV N protein staining was performed in the presence of 0.1% protease in pre-warmed phosphate buffered saline (PBS) for 10 minutes at 37° C. Antigen retrieval for other stainings was performed in citrate buffer (10 mM, pH = 6.0) with heat induction. Sections were incubated with primary antibody overnight at 4° C before incubation with secondary and tertiary antibodies. For IF assays, the slides were mounted with ProLong Diamond Antifade Mountant with DAPI (Thermo Fisher Scientific) prior to fluorescence detection with the inverted laser scanning microscope.

## Funding

BMN and TBHG were supported by Aidsfonds (P-11118), European Research Council, Advanced grant (670424). BML is supported by the Indonesian Endowment Fund for Education (grant no. 20150822023688). The funders had no role in study design, data collection and analysis, decision to publish, or preparation of the manuscript.

## Acknowledgment

We are grateful to the Boerhaave Medical Centre (Amsterdam, the Netherlands) and A. Knottenbelt (Flevo Clinic Almere, the Netherlands) for the provision of human skin tissues, Peter R. W. A. van Run and Daryl Geers for the excellent technical assistance.

**S1. MV-infected LCs were not observed after *ex vivo* infection of human epidermal sheets.** LCs (magenta) were present in abundance in human epidermal sheets. MV-infected cells (green) appeared at 2 dpi and their number increased by 4 dpi. However, none of these infected cells were LCs. Magenta: CD1a; Green: GFP; Blue: DAPI. Scale bars: 200 μm.

**S2. Nectin-4 was expressed at a relatively low level in proliferating human primary keratinocytes.** The expression level increased during differentiation. The expression of nectin-4 was abrogated in EDSS patient’s keratinocytes and differentiation did not result in increased nectin-4 expression. NCI-H358 and BLCL were included as positive and negative controls of nectin-4 expression, respectively.

